# Assessment of the Degree of Coincidence between Differentially Expressed Genes in Pancreatic Cancer with and without CAR T Cell treatment

**DOI:** 10.1101/2024.04.15.589636

**Authors:** Alibeth E. Luna-Alvear, Deiver Suárez-Gómez, Andrea A. Sanchez-Castro, Alexandra C. Rentas-Echeverria, Mauricio Cabrera-Ríos, Clara E. Isaza

## Abstract

Treatment of cancer with CAR T Cells has steadily become a viable and promising cellular therapy approach in recent years. It is well known that liquid cancers are better suited for this kind of treatment, as opposed to solid cancers. This work focuses on contrasting lists of differentially expressed genes (DEGs) found in pancreatic cancer -a solid cancer-against lists of DEGs found in post-CAR T Cell treatment of pancreatic cancer. It is postulated that the degree of coincidence in these lists could positively correlate with treatment effectiveness. OBAMA, a proprietary mathematical optimization-based analysis pipeline that minimizes user selection bias is employed here to preserve objectivity. The study utilized publicly available microarray experiments. The results indicate overall low degrees of coincidence, which partially support the postulate of this work.

## Introduction

Cancer treatment has traditionally relied on surgery, chemotherapy, and radiation. However, in 2017 the Food and Drug Administration (FDA) approved six Chimeric Antigen Receptor T-Cells (CAR T Cells) therapies. In the widest application practice of CAR T cells therapies, a patient’s own T cells are genetically engineered to express a synthetic receptor that binds specific tumor antigens (1). These modified T cells are then infused back into the patient to attack and destroy chemotherapy-resistant cancer cells. CAR T cells therapies have been proven effective in the treatment of blood cancers like leukemia and myeloma. CAR T Cells return to the bloodstream and lymphatic system and have, therefore, more contact with cancer blood cells. Nevertheless, in solid tumors CAR T Cells may not be able to penetrate tumor tissue through the vascular endothelium (2). Finding, entering, and surviving solid tumors and their microenvironments (3,4) are thus important challenges for treatment effectiveness and are the foci of current research in the field.

This work postulates the possibility that differential expression of cancer biomarker genes could be a predictor of CAR T Cell treatment effectiveness. To this end, DEGs extracted from the analysis of comparative experiments involving a particular type of cancer vs. control samples can be contrasted to DEGs resulting from the analysis of comparative experiments involving pre- and post-CAR T Cell treatment in that particular type of cancer. The degree of coincidence between these two lists of DEGs can then be correlated with the reported effectiveness of said treatment.

This study included publicly available experiments of microarrays related to pancreatic cancer (PC). Microarrays allow measuring the relative expression of thousands of genes simultaneously and the resulting datasets can subsequently be analyzed to identify DEGs. Multiple Criteria Optimization (MCO) was used to identify the DEGs in the experiments selected due to its capacity to preserve objectivity and repeatability (5). MCO can be accessed in the address (https://github.com/DeiverSuarez/OBAMA) as part of a proprietary larger tool suite called Optimization-based Analysis of Multiple Arrays (OBAMA).

The rationale to focus on PC partially obeys to the facts that it is the fourth leading cause of cancer death in the United States, and that it is projected to move up to second place by 2040(6). PC cells form solid tumors and are known to spread rapidly. Men have a higher incidence of PC, and lower survival rate than women (7,8). PC incidence in North America is reported at 8.7 per 100,00 for men, while for women it is 6.5 per 100,000, (9). Higher incidence in men has been attributed to occupational and environmental risks, alcohol and nicotine consumption, and genetic factors(10). Following from this evidence, it was important that the analysis presented here treated samples from men and women separately when feasible.

In summary, this manuscript aims to assess if CAR T Cell treatment effectiveness of cancer can be associated to the degree of coincidence between (i) DEGs identified when CAR T Cells are used and (ii) DEGs identified for the type of cancer under study. The study is centered in PC through the optimization-based analysis of comparative microarray experiments. The results are the first steps towards developing CAR T Cells treatment effectiveness predictors in cancer.

## Materials and Methods

Selection of DEGs in this work was carried out through Multiple Criteria Optimization (MCO), (5,11,12). MCO can be used to identify DEGs from the analysis of single microarray datasets as well as from the meta-analysis of multiple microarray datasets. The coded MCO tool can be found in the address, https://github.com/DeiverSuarez/OBAMA. To understand MCO in general terms, it is first important to realize that the relative change of a gene’s expression between two states (treatment vs. control) in a comparative experiment can be me measured using different criteria. With the choice of a particular criterion, it also becomes necessary to define a threshold for significance. In addition, a standardization procedure must be chosen to foster comparability, and in the presence of multiple experiments, additional transformations must be applied to deal with incommensurability. An analyst makes all these choices and, in doing so, biases the results. MCO represents each gene in an experiment or a series of experiments through multiple criteria, to then find those genes showing the best possible compromises among them. Mathematically speaking these are called the Pareto-efficient solutions, and they integrate the Pareto-efficient frontier of the multiple criteria optimization problem. The procedure does not require the analyst to decide upon any threshold values for significance and does not require standardization or transformations to deal simultaneously with experiments in different units of measurement. In this way, MCO provides a way to detect DEGs from single or multiple array comparative experiments in an objective and repeatable manner. A more detailed explanation of MCO and its code can be found in (5).

MCO is the entity selection component of the analysis pipeline called OBAMA (https://github.com/DeiverSuarez/OBAMA) which includes a correlation structure component (MST) and a biological contextualization component (OGF). Because all components of OBAMA are mathematical optimization-based, their collective nature is better described as *BioOptimatics*. Figure 1 describes the general analysis flow used to detect DEGs from microarray experiments using OBAMA and MCO This work was organized in two stages: the first one comprised the selection of suitable microarray datasets, and the second one entailed the identification of different DEG lists and from the selected datasets as well as their subsequent comparison. These stages are described next.

**Figure 1:**
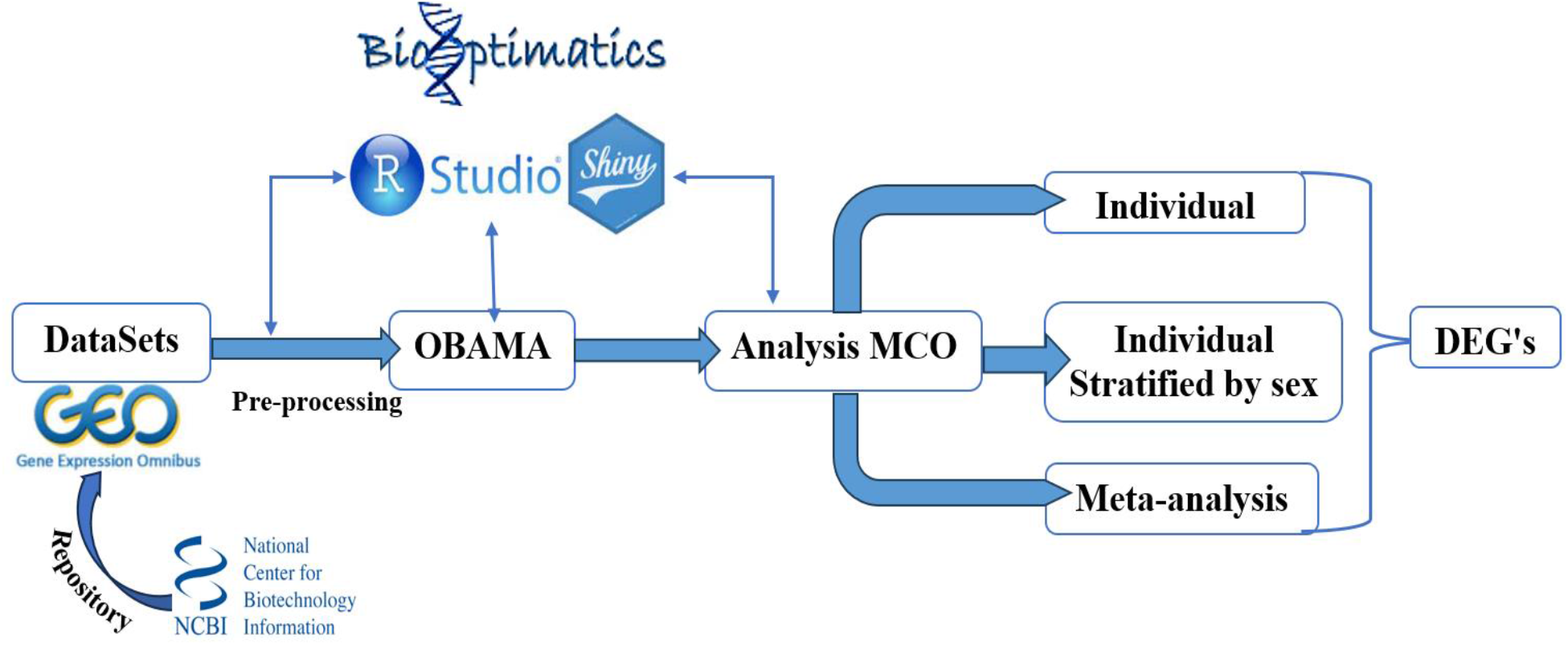
BioOptimatics workflow in this work.

### Stage1: GEO Datasets Selection

The first step was to select datasets related to PC from microarray experiments from the National Center for Biotechnology Information, specifically from Gene Expression Omnibus (GEO) (13). To be included in this study, a PC related dataset was required to contain microarray experiments organized in a comparative condition vs. control format, contain at least five samples per group of comparison, and preferably include samples from males and females. Table 1 describes the datasets selected for inclusion and their characteristics.

**Table 1.**
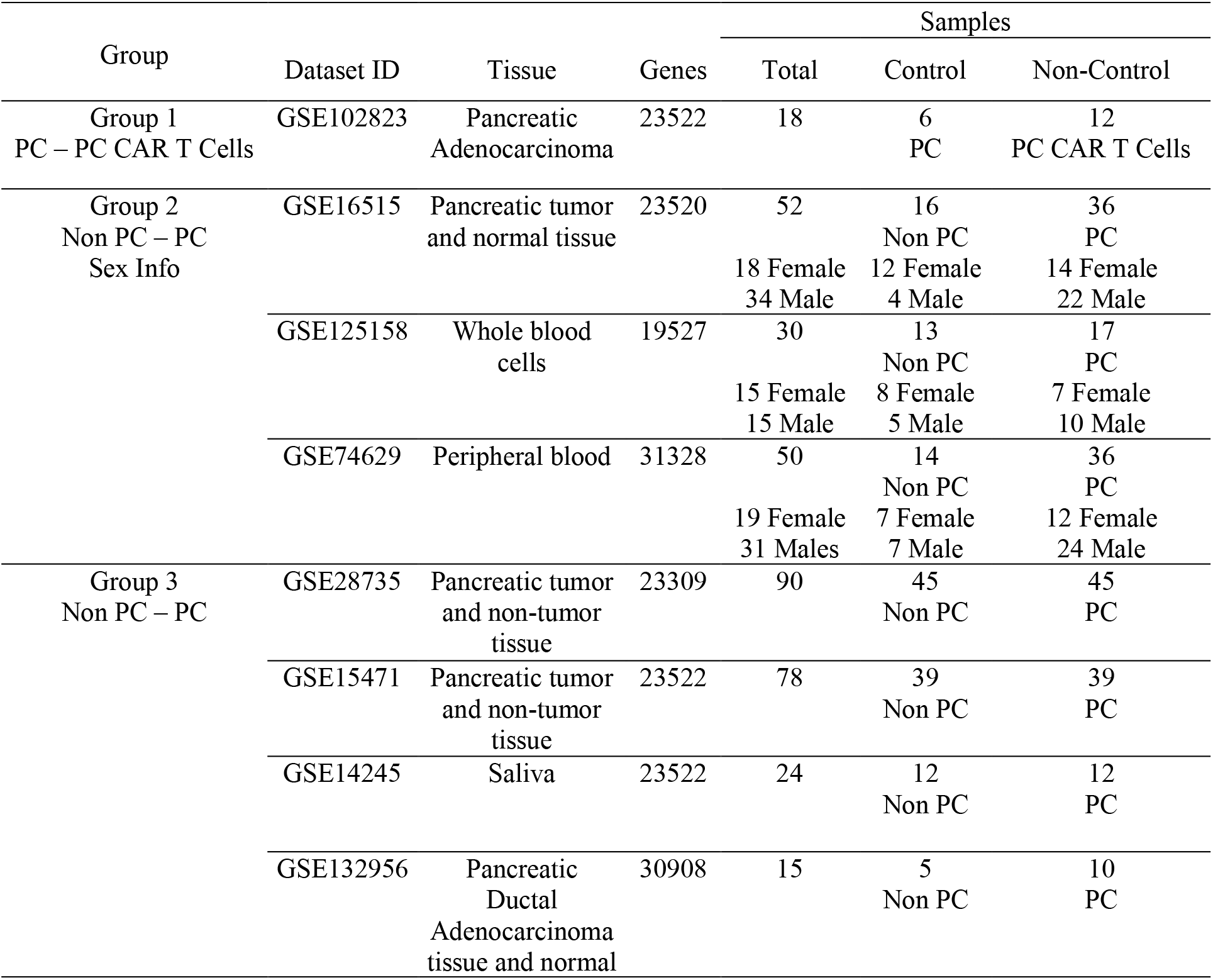
Microarray Experiments Pancreatic Cancer (PC) – Datasets’ information and groups .

Following Table 1, a total of eight microarray datasets were selected from five different GEO platforms (GPL). These were subsequently organized into three groups. Group 1 includes a single PC dataset with pre- and post-CAR-T cell treatment (GSE102823 (15)) on HPAC cell lines acquired from the American Cell Type Culture Collection (14); Group 2 has three PC datasets (Non PC vs. PC) without CAR T cell treatment and with sex information ((GSE16515, (16), (GSE125158, (17), GSE74629, (18))); and Group 3 has four PC datasets (Non PC vs. PC) without CAR T-cell treatment and without sex information ((GSE28735, (19)), (GSE15471, (20,21)), (GSE14245, (22)), (GSE132959), (23)).

### Stage 2: MCO analysis

MCO was used to conduct three types of analyses in this stage: individual dataset analysis, individual dataset analysis stratified by sex, and meta-analysis. These are summarized next for each group of datasets.

- *Individual MCO Analysis, Group 1*

This analysis characterized the effect of CAR T Cell treatment in PC. In ten Pareto efficient frontiers, thirty DEGs were identified. The DEGs in this group constituted the Reference Set against which all comparisons were made.

- *Individual MCO analysis and MCO meta-analysis, Group 2*

Individual MCO analysis stratified by sex was conducted in each of the three datasets in Group 2. The resulting DEGs were compared with the DEGs from Group 1. Subsequently, a meta-analysis involving all three datasets simultaneously was conducted (MA1) and the resulting DEGs were compared to those in Group 1.

- *Individual MCO analysis and MCO meta-analysis, Group 3*

As in the preceding task, individual MCO analyses were conducted for each dataset in group 3 to then be compared to the DEGs in group 1. A meta-analysis involving all four datasets simultaneously was also conducted (MA2), and the resulting DEGs were compared to those from the analysis of Group1.

- *Comparative analysis between MA1, MA2, and Group 1*.

This analysis involved finding the intersection of the DEG lists from MA1, MA2 and Group 1. In contrast to the previous analyses, this task was aimed to find genes that persisted and therefore could have a vital role as treatment targets.

## Results and Discussion

### MCO analysis in Group 1

This first analysis resulted in thirty DEGs. These are listed on Table 2 along with references related to their role in PC. Figure 2 helps visualize the MCO results: thirty DEGs organized in ten Pareto-efficient frontiers.

**Table 2.**
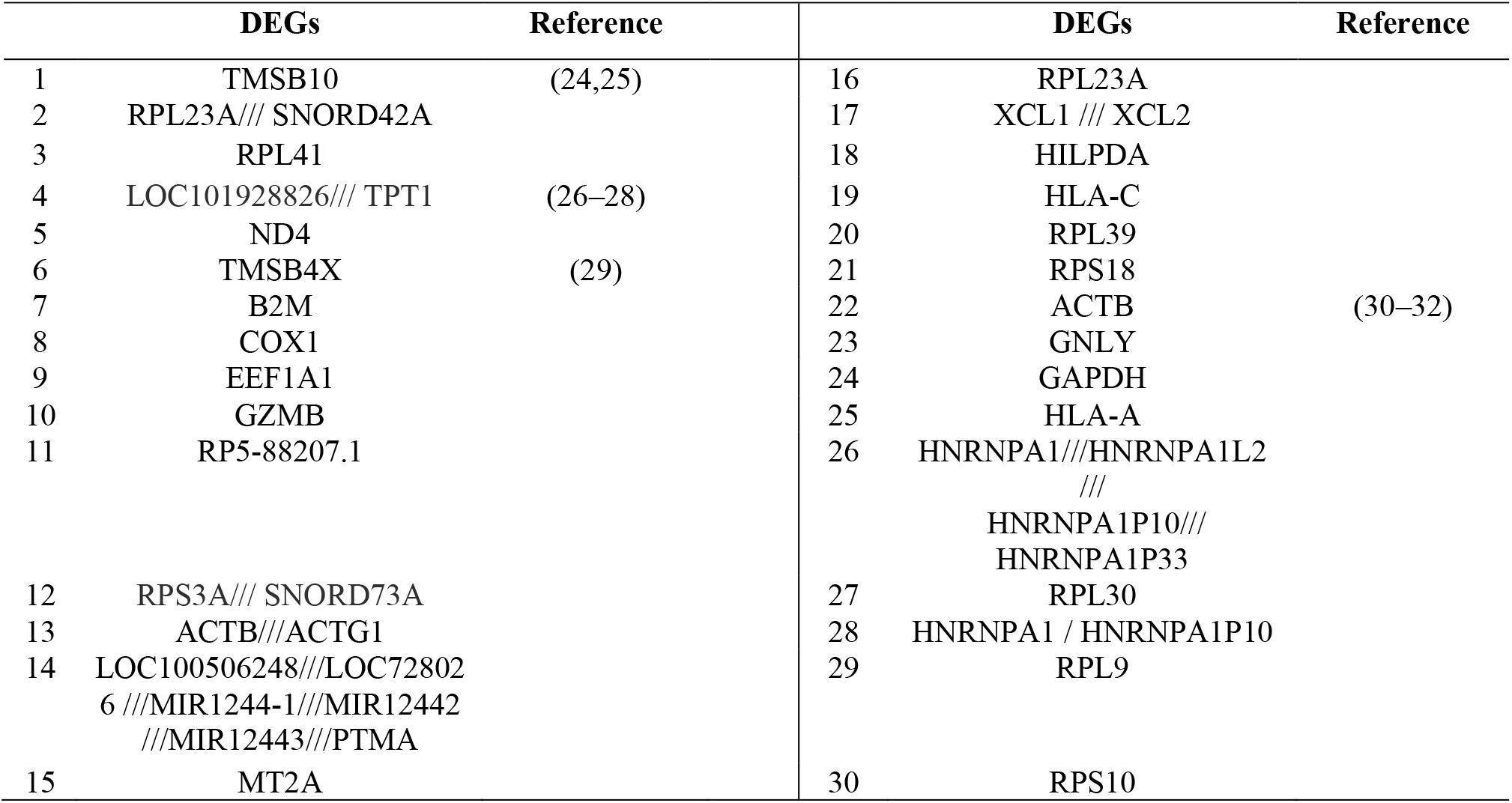
The reference set: thirty DEGs resulting from the MCO analysis of Group 1.

**Figure 2:**
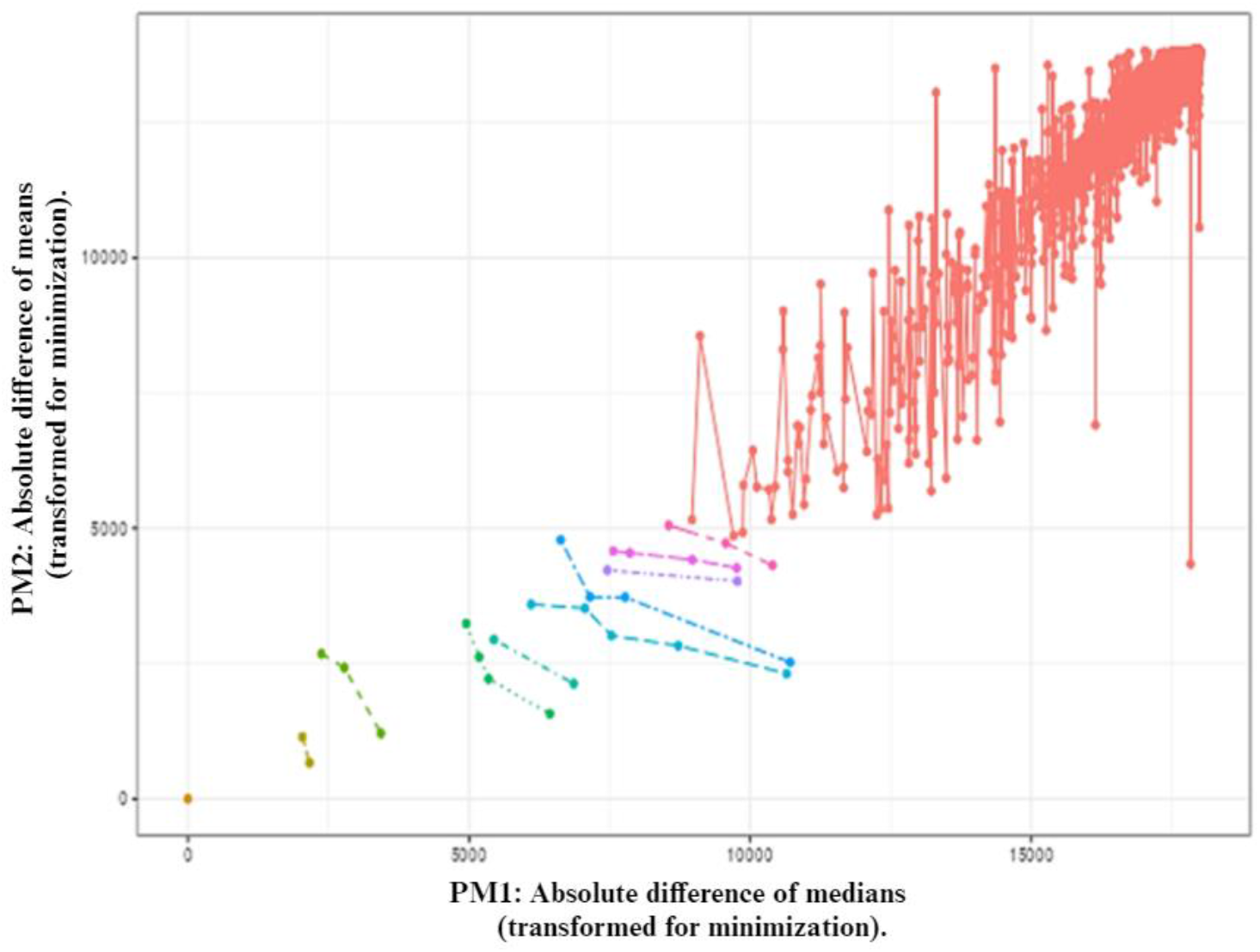
Ten efficient frontiers containing the Reference Set: 30 DEGs from the MCO analysis of Group 1.

### MCO individual analysis and MCO meta-analysis, Group 2

Table 3 shows the results of all the MCO analyses for Group 2. First, the individual analysis of each dataset resulted in DEGs that, when compared to the reference set (DEGs Group 1) resulted in low values of coincidence: 1/30 (3.33%), 0/30, and 0/30 for datasets GSE16515, GSE125158 and GSE74629, respectively. The sole coincidence occurred in gene ACTB. This gene was found to be overexpressed in GSE16515, while underexpressed in Group 1.

**Table 3.** MCO Analyses for Group 2.

| Group | Data Sets | MCO Analysis of Interest | DEGs | Coincidence with DEGs from Group 1 | DEGs in Common with Group 1 |
| --- | --- | --- | --- | --- | --- |
| Group 2<br>Non PC – PC | GSE16515 | MCO PC – Non PC | 26 | 1 / 30 | ACTB |
| (Individual datasets) | GSE125158 | MCO PC – Non PC | 62 | 0 / 30 |  |
|  | GSE74629 | MCO PC – Non PC | 48 | 0 / 30 |  |
| Group 2<br>Non PC – PC | GSE16515 | MCO Female | 24 | 1 / 30 | TMSB4X |
|  |  | MCO Male | 27 | 2 / 30 | ACTB, CTRB2 |
| (Individual datasets with sex info) | GSE125158 | MCO Female | 48 | 0/30 |  |
|  |  | MCO Male | 118 | 0/30 |  |
|  | GSE74629 | MCO Female | 39 | 0/30 |  |
|  |  | MCO Male | 30 | 0/30 |  |
| 2<br>Non PC – PC<br>MA1 | GSE16515<br>+<br>GSE125158<br>+<br>GSE74629 | MA1 | 496 | 9 / 30 | TMSB10, RPL39,<br>TMSB4X, ACTB,<br>B2M, EEF1A1,<br>MT2A, RPL23A,<br>RPS18 |

Furthermore, because these datasets have information regarding the sex of the patients and compare PC samples with control samples, it is possible to have four subgroups on each dataset: female-control, female-PC, male-control, male-PC. Figure 3a shows these four subgroups and how MCO was applied to contrast control subgroups (MCO Control), PC subgroups (MCO PC), Female subgroups (MCO Female), and Male subgroups (MCO Male). Figure 3b shows Venn diagrams displaying the number of resulting DEGs per MCO application for each of the three datasets separately. In this work, the focus was on MCO Female and MCO Male, which generated lists of PC DEGs per sex. When the PC DEGs per sex were compared to the list of 30 DEGs from Group 1, there were few coincidences, which occurred only when using dataset GSE16515: 1/30 and 2/30 for MCO Female and MCO male, respectively. No other coincidences were found using the rest of the datasets.

**Figure 3.**
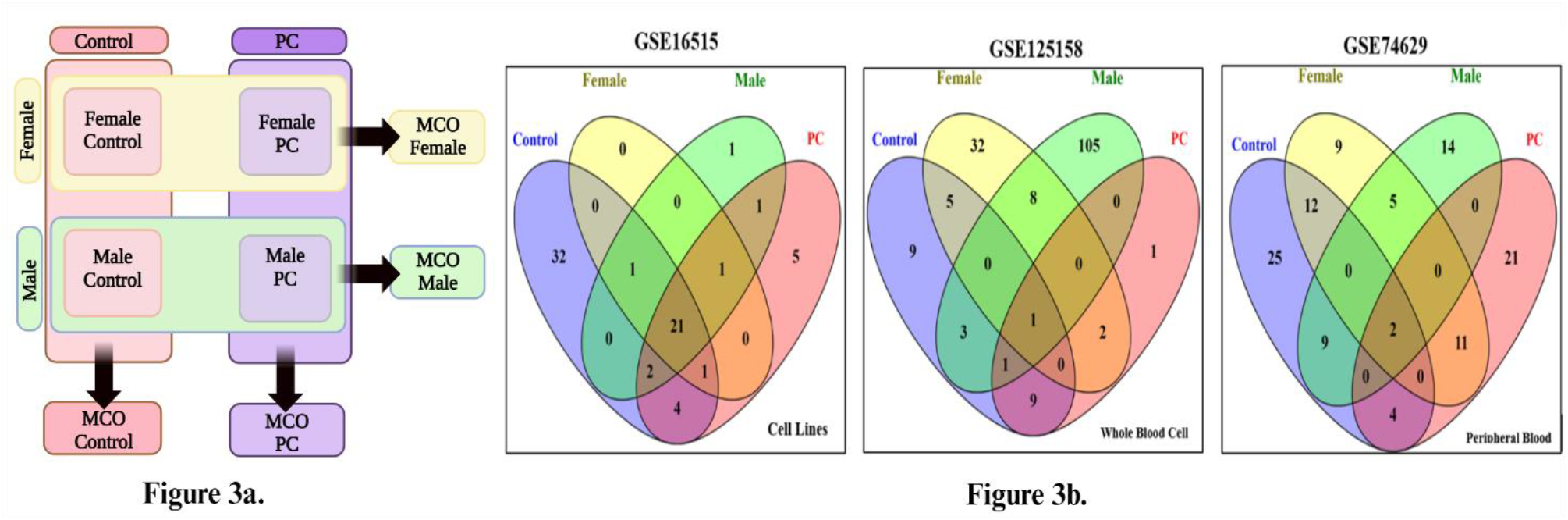
Figure 3a. MCO analyses in four subgroups; Figure 3b Venn-Euler diagrams of MCO results in four subgroups for datasets GSE16515, GSE125158 and GSE74629.

TMSB4X was the only coincidence between the DEGs from MCO Female (dataset GSE16515) and the reference set. It was found to be overexpressed both in GSE16515 and in the reference set from Group 1. On the other hand, ACTB and CTRB2 were the two coincidences between DEGs from MCO Male (dataset GSE16515) and the reference set. Both genes were found to be overexpressed in GSE16515 and in the reference set.

MA1 included the simultaneous analysis of the three datasets in Group 2. At five efficient frontiers, this analysis converged to 496 DEGs. When these were compared to the reference set, there was a coincidence of 9/30 genes (30%). The list of genes is {TMSB10, RPL39, TMSB4X, ACTB, B2M, EEF1A1, MT2A, RPL23A, RPS18}. The relative expression changes of these genes are shown in Table 5.

### MCO individual analysis and MCO meta-analysis, group 3

Table 4 shows the results of all the MCO analyses for Group 3. These were also at the individual dataset level as well as at the meta-analysis (MA2) level. The resulting DEGs, however, did not have a single coincidence with the reference set for the datasets GSE28735, GSE15471, and GSE132956 individually. The only coincidence occurred in dataset GSE14245 for a level of 1/30 (3.33%). This coincidence occurred in gene LOC101928826///TPT1. This gene was found underexpressed in dataset GSE14245, while overexpressed in the reference set (see Table 5).

**Table 4.**
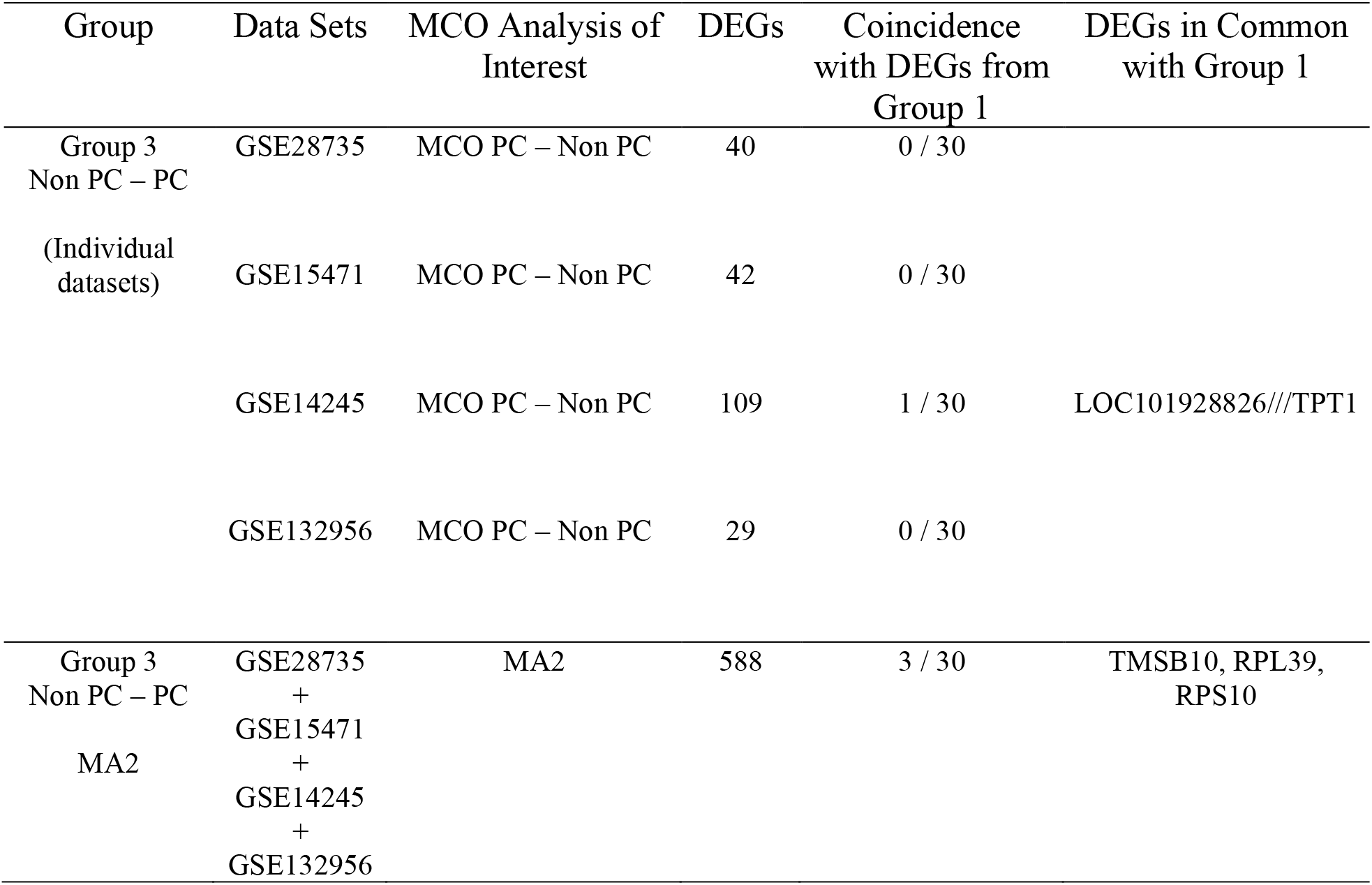
MCO Individual Analysis for group 3.

MA2, at five efficient frontiers, identified 588 DEGs. In a comparison with the reference set, three genes were found in common: TMSB10, RPL39 and RPS10; for a coincidence level of 3/30 (10%). Changes in expression can be consulted in Table 5.

**Table 5.**
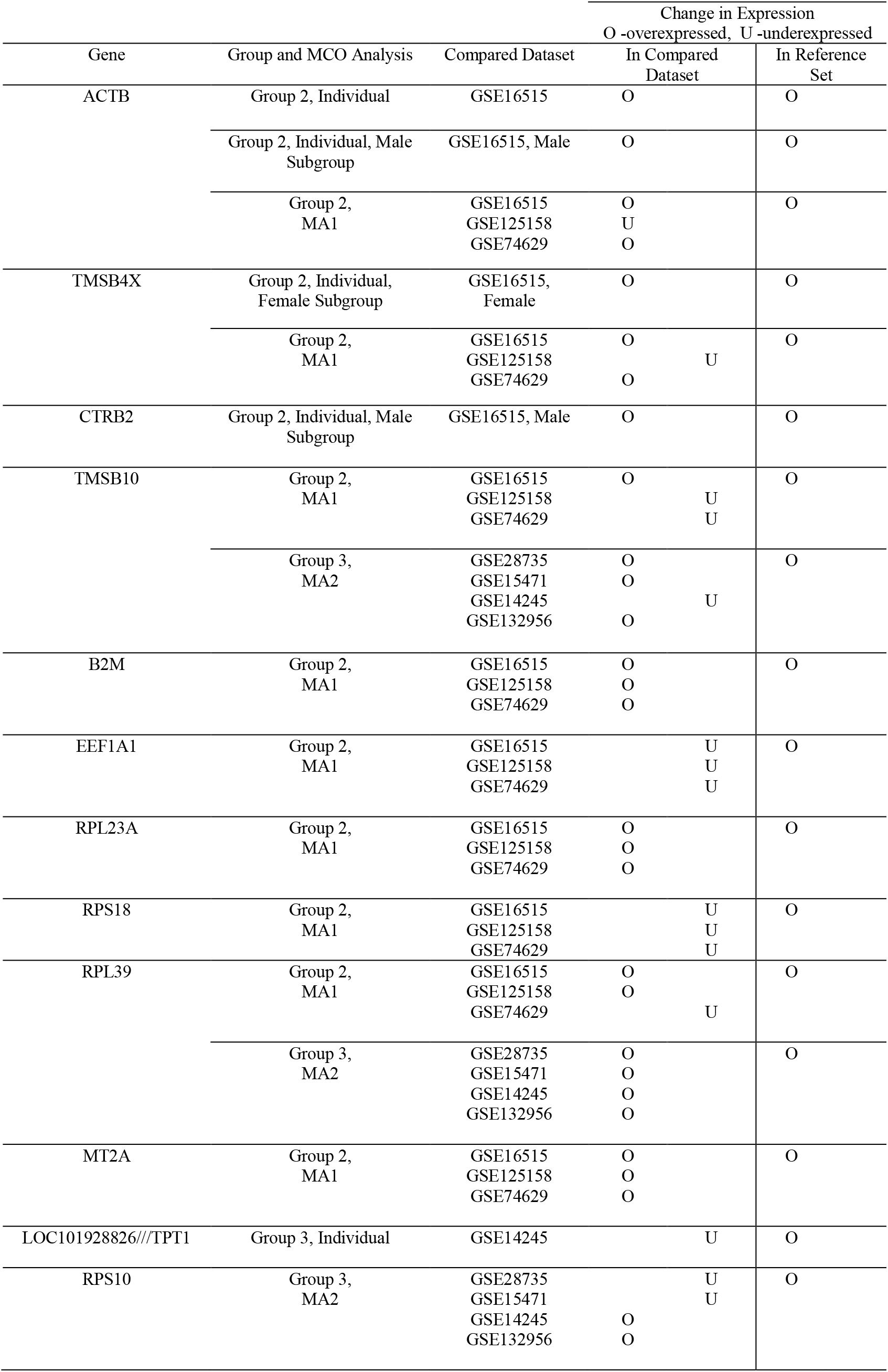
Expression changes in the genes that coincided with the reference set on each analysis.

### Comparative analysis between MA1&MA2, and MPC

An intersectional analysis was conducted between the results from MA1 and MA2 resulting in 85 genes. These genes were then compared to the reference set resulting in two matching genes: TMBS10 and RPL39. Both were found to be overexpressed after CAR T Cell treatment of PC in Group 1 (Table 5). Because these genes persist throughout most analyses in this work, information on them is provided in Appendix A, as they can have potential as treatment targets.

## Conclusion

This work postulated the possibility that effectiveness of CAR T Cell treatment in cancer could be related to the coincidence level between DEGs from (i) pre- and post-CAR T Cell treatment samples and (ii) control and cancer samples. To this end, a BioOptimatics study in pancreatic cancer was conducted using publicly available microarray datasets. The resulting coincidence levels ranged from zero to a maximum of nine genes in common. These numbers are deemed low when compared to a reference set of 30 DEGs found in a dataset contrasting pre- and post-CAR T Cell treatment of pancreatic cancer as a reference. CAR T Cells have been considered less effective in solid tumor cancers, like the one studied here, so in this regard the results partially support the premise of this work. A complete assessment will come from the future study of the application of CAR T Cells to blood cancers in a manner similar to this work.

## Abbreviations

*PC*: Pancreatic Cancer
*CAR*: Chimeric antigen receptor
*MCO*: Multiple Criteria Optimization
*NCBI*: National Center for Biotechnology Information
*GEO*: Gen Expression Omnibus
*GPL*: Platform Accession Number
*OBAMA*: Optimization-Based Analysis of Multiple Arrays
*DEG*: Differentially expressed genes.
*MA*: Meta-analysis
*NCI*: National Cancer Institute

## Acknowledgements

This project was supported by the NSF Engineering Research Center for Cell Manufacturing Technologies (CMaT)

## Author Contributions

### Conceptualization

Alibeth Luna, Mauricio Cabrera-Ríos, Clara E. Isaza.

### Data curation

Alibeth Luna-Alvear, Andrea A. Sánchez-Castro, Alexandra C. Rentas-Echeverria

### Formal analysis

Alibeth Luna, Mauricio Cabrera-Ríos, Clara E. Isaza.

### Investigation

Alibeth Luna-Alvear, Andrea A. Sánchez-Castro, Alexandra C. Rentas-Echeverria, Mauricio Cabrera-Ríos, Clara E. Isaza.

### Methodology

Alibeth Luna-Alvear, Deiver Suárez-Gómez, Mauricio Cabrera-Ríos, Clara E. Isaza.

### Project administrations

Mauricio Cabrera-Ríos, Clara E. Isaza.

### Writing original draft

Alibeth E. Luna-Alvear

### Writing review & editing

Alibeth E. Luna-Alvear, Mauricio Cabrera-Ríos, Clara E. Isaza.

## APPENDIX A TMSB10

Thymosin β10 (TMSB10) is a member of the thymosin family and plays a key role in various physiological processes such as tissue regeneration and inflammatory regulation. In addition, it has been observed that its expression is increased in several types of carcinomas. TMSB10 has been found to be overexpressed in several types of human cancer including renal cell carcinoma (33), non-small cell lung cancer, and papillary thyroid cancer specifically. On the other hand, in (34), it was demonstrated that high expression of TMSB10 could serve as a useful diagnostic and prognostic biomarker and a potential therapeutic target for clear cell renal cell carcinoma. Consistent with (35), upregulation of TMSB10 has been observed in breast cancer cells and tissues, suggesting that it could be a promising minimally invasive serum cancer biomarker for breast cancer diagnosis. Relative to PC, the authors of (24) found that TMSB10 expression was upregulated in both human pancreatic carcinoma tissues and cell lines, and suggested a potential role of TMSB10 in the carcinogenesis of pancreatic carcinoma. TMBS10 was also found to be one of the characteristic genes associated with pancreatic cancer in (25). TMBS10 is overexpressed even after CAR T cell treatment for the data used in this work.

RPL39

Ribosomal protein L39 is a protein coding gene which has been shown to play a role in several types of cancers. RPL39 in pancreatic cancer is linked to cancer cell regression and apoptosis enhancement. This gene is overexpressed in the most aggressive pancreatic cancer cell lines PANC-1 and MIA PaCa-2. Because of the effect of RPL39 on pancreatic cancer cell apoptosis, targeting RPL39 could be a potential treatment in pancreatic cancer (36). The RPL39 gene has been shown to be up-regulated after long-term silencing of oncogenic KRAS in pancreatic cancer PANC-1 cells, (37). Knockdown of RPL39 expression by RPL39-siRNA significantly inhibits the growth of human pancreatic cancer cells in vitro and in vivo. This evidence suggests that RPL39 is a potential therapeutic target for pancreatic cancer, (37). In general, ribosomal proteins play a role in the development and progression of pancreatic cancer. RPL34, RPS15A, RPL26, RPL29, RPS8, RPL15, and RPL21 have all been shown to have an effect on pancreatic cancer (36). Aside from that, RPL39 has been shown to be overexpressed in breast cancer and in hepatocellular carcinoma, which is the most common type of liver cancer, (36). RPL39L is a paralog RPL39 present in mammals, and it is exclusively expressed in male gonads (38). In the results of this work, RPL39 keeps being overexpressed after CAR T cell treatment.

